# Orientation to polarized light in tethered flying honeybees

**DOI:** 10.1101/803809

**Authors:** Norihiro Kobayashi, Ryuichi Okada, Midori Sakura

**Author notes:** Correspondence to Midori Sakura.

## Abstract

Behavioral responses of honeybees to a zenithal polarized light stimulus were observed using a tethered animal in a flight simulator. Flight direction of the bee was recorded by monitoring the horizontal movement of its abdomen, which was strongly anti-correlated with its torque. When the e-vector orientation of the polarized light was rotated clockwise or counterclockwise, the bee responded with periodic right-and-left abdominal movements; however, the bee did not show any clear periodic movement under the static e-vector or depolarized stimulus. The steering frequency of the bee was well coordinated with the e-vector rotation frequency of the stimulus, indicating that the flying bee oriented itself to a certain e-vector orientation, i.e., exhibited polarotaxis. The percentage of bees exhibiting clear polarotaxis was much smaller under the fast stimulus (3.6 ° s^-1^) compared with that of the slow stimulus (0.9 or 1.8 ° s^-1^). The bee did not demonstrate any polarotactic behavior after the dorsal rim region of its eyes, which mediates insect polarization vision in general, was bilaterally covered with black paint. The bees demonstrated a high preference for e-vector orientations between 120 to 180°. Each bee exhibited similar e-vector preferences under clockwise and counterclockwise stimuli, indicating that each bee has its own e-vector preference, which probably depends on the bee’s previous foraging experience. Our results strongly suggest that the flying honeybees utilize the e-vector information from the skylight to deduce their heading orientation for navigation.

**Summary statement:** Tethered flying bees exhibited polarotaxis under a zenithal rotating e-vector stimulus, in which their right-and-left abdominal movements were coincident with the rotation of the stimulus.

## INTRODUCTION

As a result of sunlight scattering in the atmosphere, the skylight is partially plane-polarized and the celestial e-vectors are arranged in a concentric pattern around the sun (Strutt, 1871; Wehner, 1997). It is well known that many insects exploit this skylight polarization for visual compass orientation and/or navigation (for review: Wehner, 1994; Wehner and Labhart, 2006; Heinze, 2014). There have been an enormous number of studies about insect polarization vision, not only at the behavioral level (e.g., Dacke et al., 2003; Reppert et al., 2004; Henze and Labhart, 2007; Weir and Dickinson, 2012), but also at the neural network level, such as sensory (e.g., Blum and Labhart, 2000; Weir et al., 2016) and central brain mechanisms (e.g., Labhart 1988; Heinze and Homberg, 2007; Sakura et al., 2008; Heinze and Reppert, 2011; Bech et al., 2014). The e-vector detection in insects is mediated by a group of specialized ommatidia located in the most dorsal part of the compound eye, the dorsal rim area (DRA), in which the photoreceptors are monochromatic and highly polarization-sensitive (for review: Labhart and Meyer, 1999; Wehner and Labhart, 2006). The neural pathway of polarization vision in the brain has been documented in several species, and the central complex, one of the higher centers of the insect brain, is considered to be the location of an internal compass (for review: Homberg et al., 2011; Heinze, 2017), although it is still unclear how the central complex controls the animal’s steering during navigation.

Foraging behavior in social insects, such as ants and bees, is a useful model system for studying insect navigation because they repeatedly forage back and forth between the nest and a feeding site. In particular, the path integration mechanisms in the desert ants *Cataglyphis* have been extensively studied in regard to insect navigation (Wehner, 2003; Collett and Carde, 2014), and *Cataglyphis* is well known to choose their heading direction using celestial polarization cues during long distance navigation (Fent, 1986; Wehner, 1997; Wehner and Müller, 2006). In addition to path integration based on the polarization compass, ants could learn visual landmarks or panoramic views at familiar locations and use them for local navigation (Collett et al., 1992; Wehner et al., 1996; Collett et al., 1998; Graham and Cheng, 2009; Narendra et al., 2013). Honeybees also undertake long-distance foraging trips, that may reach over 40 km (Couvillon et al., 2014). The foragers, after returning from a food source, transfer the food location to their nestmates by the waggle dance, in which the vector information from their nest to the food location is encoded. This clearly indicates that honeybees navigate based on path integration (Frisch, 1967). The waggle dance has also been used to clarify the utilization of polarization vision in their navigation. Under unpolarized light, the dancing honeybees failed to transfer the correct directional information to the food source and their waggle dance orientations were strongly affected by artificially-polarized light (Frisch, 1967; Rossel and Wehner, 1987; Sherman and Visscher, 2002). More recently, it was reported that they changed their waggle dance orientations depending on the e-vectors of polarized light experienced during their foraging trip (Evangelista et al., 2014). These results suggest that the honeybees use polarization vision for path integration, i.e., deducing the direction to the food source. In contrast to investigations regarding path integration into the waggle dance, to our knowledge, there has been only one study that examined whether a flying bee can use celestial e-vector information to choose its flight route. Kraft et al. (2011) showed that bees trained to fly in a four-armed maze to a feeder, in which the bees received polarized light stimulus from above, chose their foraging routes as they received e-vector information, similar to what they experienced during the training. This suggests that the flying bees actually sense the e-vector orientation from the sky and use it for navigation.

Similar to ants, honeybees display both path integration and visual landmark navigation. Bees use familiar landmarks to find the correct location (Cartwright and Collett, 1983; Fry and Wehner, 2005), and in some cases, visual landmarks dominate path integration (Chittka and Kunze, 1995). Furthermore, a single bee can memorize several food locations simultaneously based on the landmarks at each location and visit the best one among the destinations in a context-dependent manner (Collett and Kelber, 1988; Zhang et al., 2006). To accomplish those complicated navigational tasks, foraging bees must use multiple navigational strategies, as the situation demands, by using several visual cues such as polarized light and landmarks. However, it still remains largely unknown how honeybees behaviorally select and use polarized light or landmark cues during navigational flight. This is probably because of the technical difficulties of presenting artificial visual stimuli to freely flying bees (Evangelista et al., 2014).

Recently, some behavioral studies have overcome this difficulty by using tethered bees (Luu et al., 2011; Taylor et al., 2013), with which we can observe responses in flying behavior of the dorsally tethered bees to lateral optic flow and frontal air-flow stimuli. It was observed that the tethered bees showed “streamlining” responses, whereby they raised their abdomen in a correlated manner with the speed of the optic and air-flow stimuli. In the present study, we investigated how the flying bees respond to polarized light stimuli using the tethered system. We constructed a flight simulator, in which we could examine the tethered bee’s flight response to a rotating polarized stimulus and found that they tended to orient themselves to a certain e-vector direction, i.e., they exhibited clear “polarotaxis,” during the flight.

## MATERIALS AND METHODS

### Animals

The honeybees, *Apis mellifera L.*, used in this study were reared in normal ten-frame hives on the campus of Kobe University. Forager honeybees with pollen loads were collected at the hive entrance before the experiment and anesthetized on ice or in a refrigerator. An L-shaped metal rod for tethering was attached to the pronotum of an anesthetized bee, as previously described (Luu et al., 2011). Briefly, the hair on the pronotum was gently shaved using a small piece of a razor blade, and the metal rod was adhered using a small amount of light-curing adhesive (Loctite; Henkel, Dusseldorf, Germany). The following image analyses of bee behavior (see below) were conducted by marking the tip of the abdomen with a white light-curing dental sealant (Conseal f; SDI Ltd., Bayswater, Australia). Next, the bees were placed in a warm room to recover from anesthesia and fed several drops of 30 % sucrose solution before the experiment.

### Setup

The experiments were performed using a custom-made black box (Fig. 1) in a dark room. A tethered bee was mounted in the box by attaching the end of the metal rod to a three-dimensional manipulator such that the bee’s location could be adjusted manually. The flying behavior of the tethered bee was enhanced by stimulating the bee with a headwind from an air circulator and optic flow from a PC monitor. The circulator was located outside of the box and connected to a tunnel that carried the wind stimulus into the box. The end of the tunnel (diameter, 8 cm), consisting of many fine plastic straws to reduce the turbulent flow of wind, was fixed at 10 cm from the bee’s head. The wind speed was 1.7–2.0 m s^-1^. The PC monitor (RDT1711LM; Mitsubishi Electric, Tokyo, Japan), covered with a sheet of tracing paper to eliminate any polarized components of the light, was located 5 cm beneath the tethered bee. The optic flow stimulus of moving black-and-white stripes was displayed on the monitor using a self-made program in Microsoft Visual C^++^. The spacing of the stripes and the speed of the stimulus as seen by the bee were approximately 40 ° s^-1^ and 900 ° s^-1^, respectively.

**Fig. 1.**
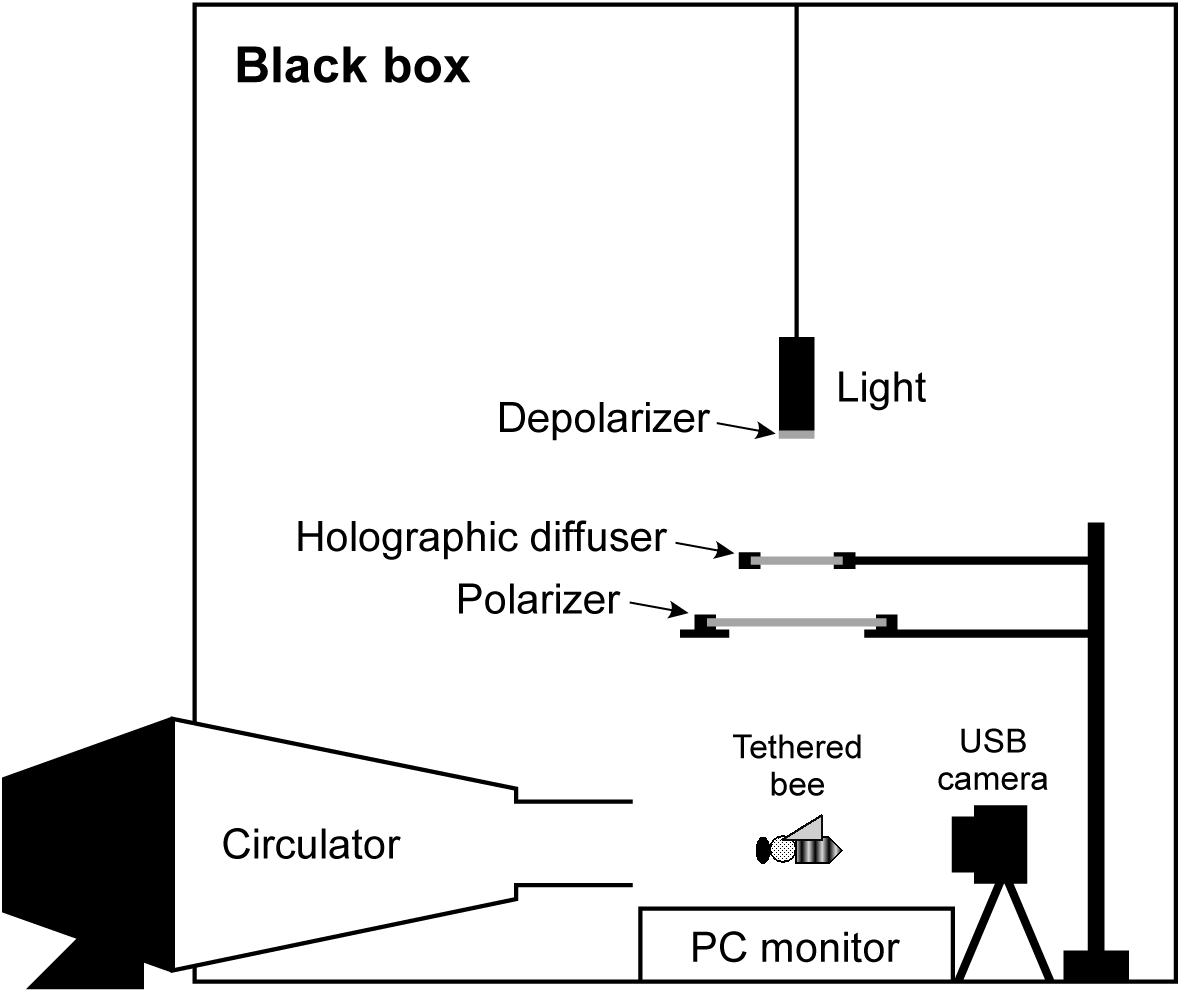
Experimental setup. Light from a xenon lamp was equally depolarized and then linearly polarized using a UV-transmitted polarizer. A bee was tethered under the polarizer and its flight was monitored by a USB camera. For the stable flight of a tethered bee, rectified wind from a circulator and moving black-and-white stripes on a PC monitor were presented.

Light from a xenon lamp (LC8; Hamamatsu Photonics, Hamamatsu, Japan) was applied above the bee using a quartz light guide. The light was filtered using a depolarizer (DPU-25; ThorLabs, Newton, NJ) at the end of the light guide to eliminate any polarized components of the light and a holographic diffuser (48-522; Edmund Optics, Barrington, NJ) was clamped under the end of the light guide. The diffuser reduced illuminance irregularity and increased the size of the light fit around a linear polarizer (HN42HE; diameter, 15 cm; Polaroid Company, Cambridge, MA) beneath the diffuser. The polarizer was mounted on a circular holder that could be rotated using a DC motor. The stimulus was centered at the bee’s zenith (with respect to flying head position) at a distance of 15 cm, providing a dorsal, polarized stimulus of 53 ° in diameter. In the experiments, in which unpolarized light stimulus was used, the depolarizer was clamped just above the bee’s head instead of at the end of the light guide such that the size of the light stimulus covered the entire receptive field of the bee’s DRA. The intensity of the polarized and unpolarized white light at the animal level was approximately 1000 lx.

### Behavioral experiments

A bee with the metal rod was fixed in the experimental box after complete recovery from anesthesia. First, we let the bee hold a small piece of paper so that it could not start flying. The e-vector angle of the polarizer was set at 0° with respect to the bee’s body axis, and static black-and-white stripes were displayed on the PC monitor. After the bee had been familiarized with the box, the paper was removed to allow the bee to start flying, and the wind and optic flow stimuli were simultaneously presented. After the bee’s flight became stable, the polarizer started rotating slowly (0.9, 1.8, or 3.6 ° s^-1^), and the behavior of the bee was monitored for 600 s. When a bee stopped flying before 600 s, the data were not used in the analysis. In some cases, the bee was tested 3 times under different stimulus conditions—clockwise (CW), static, and counterclockwise (CCW). The order of these three stimuli was randomly changed for each experiment. In other cases, a bee was tested only with the CW stimulus.

The flying behavior of the tethered bee was monitored using a USB camera (IUC-300CK2; Trinity Inc., Gunma, Japan) placed behind the bee (see Fig. 1). Images of the bee were recorded at a rate of 1 Hz, i.e., 600 images for 10 min data. For each image, the x-coordinate of the bee’s abdominal tip was determined manually to estimate flying orientation (see Fig. S1). A series of x-coordinates was then calibrated into actual distances (in mm) from the center, where the tethering wire was fixed and used for further analysis (see below).

Whether the DRA of the compound eye was involved in flying behavior under the polarized light stimulus was determined using bees whose DRAs were painted (Fig. 7C, D). The DRAs were painted as in our previous work (Sakura et al., 2012) with black acrylic emulsion paint (Herbol; Cologne, Germany) under a dissecting microscope just before the tethering procedure described above. The DRA of a compound eye is visually identifiable because the cornea appears slightly grey and cloudy (Meyer and Labhart, 1981). Because it was technically not possible to cover the DRA alone, which consists of only 4–5 horizontal rows of ommatidia (see Meyer and Labhart, 1981; Wehner and Strasser, 1985), a small area of the unspecialized dorsal region next to the DRA was also painted. After the experiments, the paint cover was checked in all the experimental animals under a dissecting microscope. Data for cases in which any of the paint was missing were excluded from further analysis. The three ocelli, which are not involved in polarization vision (Rossel and Wehner, 1984), were not painted in the experiments.

### Analysis and statistics

All data analyses were performed using self-made programs in MATLAB (MathWorks Inc., MA, USA). Periodicity of the time course of the abdominal tip location was analyzed using the fast Fourier transform (FFT). For FFT, data for only the last 400 s of each trajectory (600 s in total) were used because the periodicity of a bee’s flight was occasionally unstable at the beginning of the stimulus (e.g., see Fig. 2Ac). The relative power spectrum was calculated, and peak frequencies were determined. We defined a bee to be aligned with a certain e-vector orientation or showing “polarotaxis”, when the power spectrum of the bee showed a peak at the stimulus frequency, i.e., 0.5, 0.01, and 0.02 Hz for 0.9, 1.8, and 3.6 ° s^-1^ stimuli, respectively. Distributions of bees showing polarotaxis were statistically analyzed using Fisher’s exact test or Cochran’s *Q*‒test with post-hoc McNemar test for among-or within-group comparisons, respectively. In addition, the largest peak in the power spectrum of each bee was determined to compare the distribution of the peaks by a bee.

In the case of experiments where 1.8 ° s^-1^ CW stimulus was used, a preferred e-vector orientation (PEO) for each bee that demonstrated polarotaxis was obtained from a phase of the stimulus frequency component (0.01 Hz) in the division signal after FFT. The uniformity of the distribution of PEOs was statistically analyzed using the Rayleigh test (Batschelet, 1981). For the bees showing polarotaxis to both CW and CCW stimuli, differences in the PEOs between these two stimuli were also calculated for each bee by subtracting the value of the CW stimulus from that of CCW stimulus, and the distribution of the differences was analyzed using the *V*-test with 0 ° as an expected mean angle (Batschelet, 1981). All circular statistics were performed using Oriana software (ver. 3.12; Kovach Computing Services, UK).

## RESULTS

### Polarotactic behavior of tethered bees

Under our experimental condition, approximately two-thirds of the experimental tethered bees could stably fly for over 10 min. A representative horizontal trajectory of a bee’s abdominal tip under the three different polarized light conditions is shown in Fig. 2A. When the e-vector of the polarized light stimulus was gradually (1.8 ° s^-1^) rotated clockwise or counterclockwise, the bee showed a periodic right-and-left abdominal movement, regardless of the rotational direction (Fig. 2Aa, c). The FFT analysis of the last 400 s of the trajectory data clearly showed that these abdominal movements were synchronized with an e-vector rotating frequency of 0.01 Hz (Fig. 2Ba, c). Conversely, a bee did not show such periodic movement under the static e-vector stimulus (0 ° with respect to the body axis; Fig. 2Ab), and the peak of the power spectrum (PS) was detected at 0.0025 Hz instead of at 0.01 Hz, which is coincident with the entire data length (Fig. 2Bb). In total, over half of the experimental bees showed a clear peak at 0.01 Hz in the PS under the rotating e-vector stimulus (12 and 14 of 21 bees for CW and CCW, respectively); however, under the static 0 ° e-vector stimulus, only 2 of the 21 bees showed a 0.01 Hz peak in the PS, which was significantly smaller than the number of bees showing a peak at 0.01 Hz under the rotating stimulus (data not shown; CW: *p* = 0.008, CCW: *p* = 0.001, Cockran’s *Q*-test with post-hoc McNemar test). In addition, a significantly higher number of bees (7 and 6 of 21 bees for CW and CCW, respectively) displayed the highest peaks at 0.01 Hz in the PS compared with that (none of the 21 bees) under the static 0 ° e-vector stimulus (Fig. 3; CW: *p* = 0.008, CCW: *p* = 0.014, Cockran’s *Q*-test with post-hoc McNemar test). In the averaged PS, a clear peak was noted at 0.01 Hz under the CW or CCW stimulus, although another strong peak was detected at 0.0025 Hz (Fig. 3A, C), and the peak was only detected at 0.0025 Hz under the static stimulus (Fig. 3B).

**Fig. 2.**
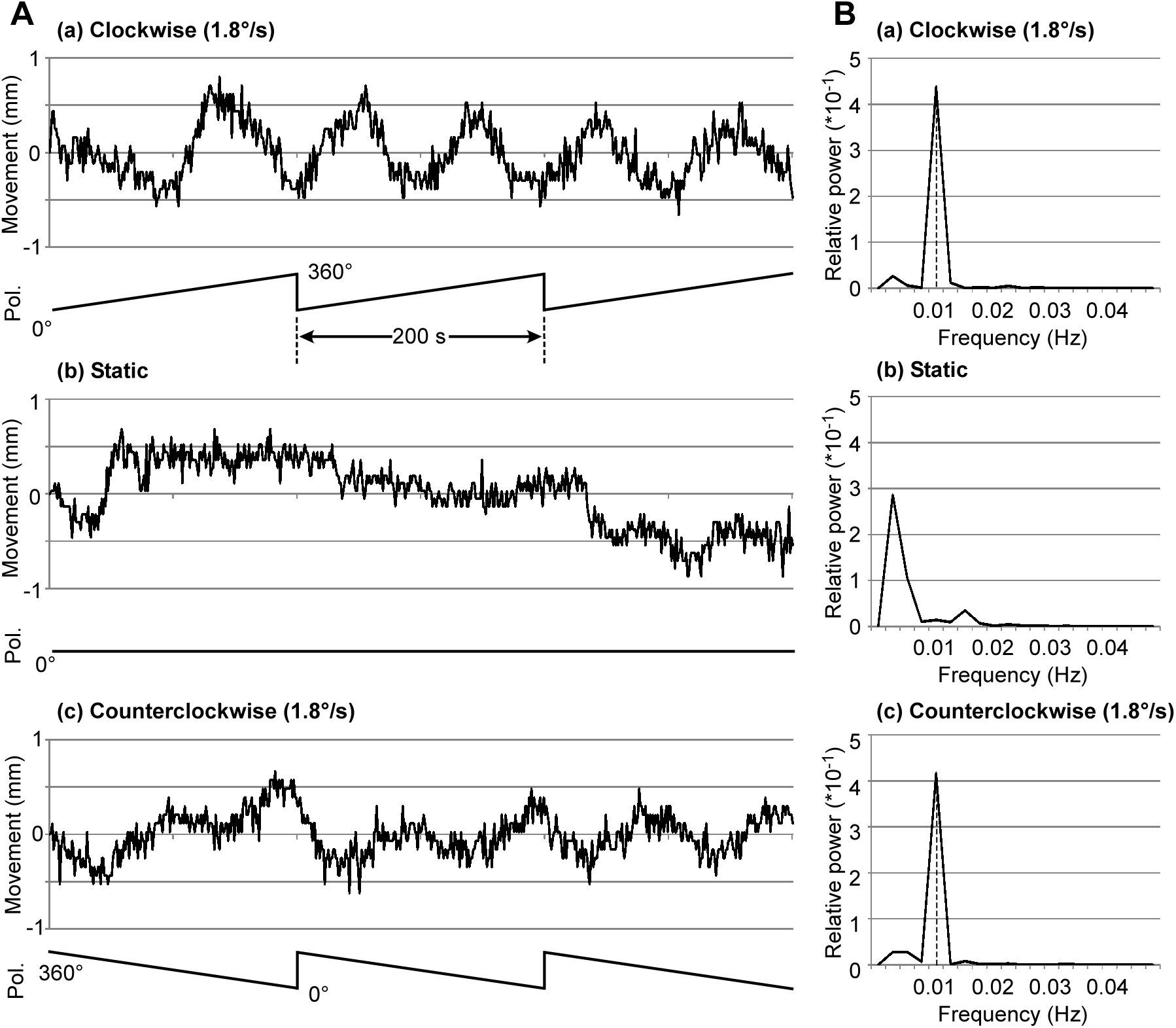
A bee’s abdominal movement under the polarized light stimulus. Trajectories of the abdominal tip (**A**) and the power spectrum (**B**) under the clockwise (1.8 ° s^-1^; **a**), static (**b**), and counterclockwise (1.8 ° s^-1^; **c**) stimulus. The lower trace in each trajectory (Pol.) indicates the e-vector orientation of the polarizer with respect to the bee’s body axis. Under rotating e-vector (**a** and **c**), the abdomen showed periodical movements from side to side. Dashed lines indicate the peaks at the stimulus rotation frequency (0.01 Hz).

**Fig. 3.**
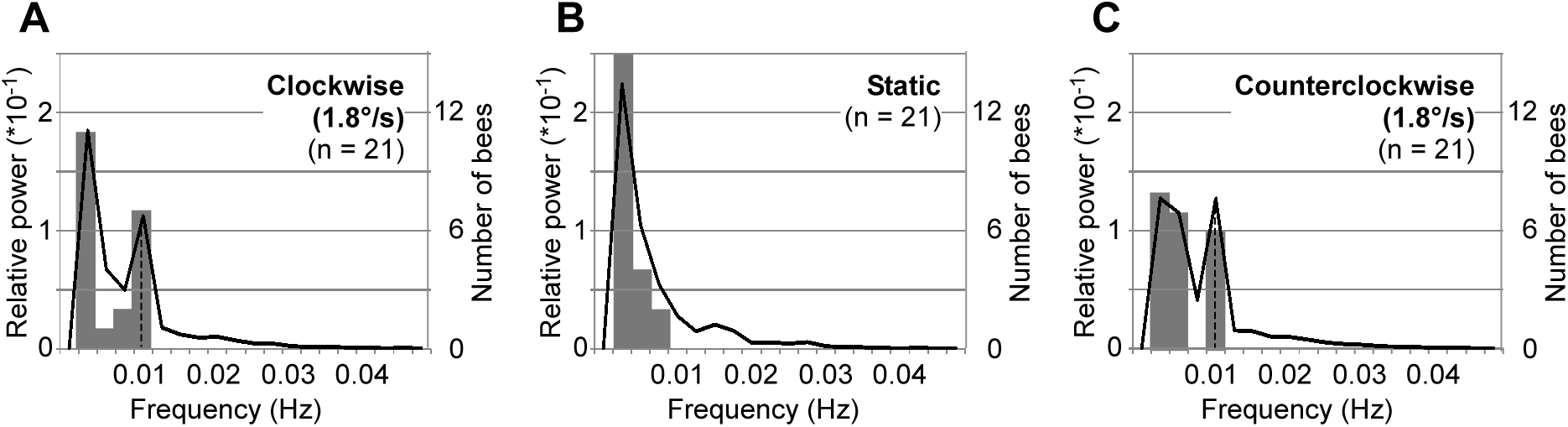
Averaged power spectrum of the abdominal movements under the polarized light stimulus. Power spectrum curves (black line) and histograms of the maximum peak in each power spectrum (gray bars) under the clockwise (1.8 ° s^-1^; **A**), static (**B**), and counterclockwise (1.8 ° s^-1^; **C**) stimulus are shown (N = 21). Dashed lines indicate the peaks at the stimulus rotation frequency (0.01 Hz).

To determine whether the periodic movements were not elicited by the rotation of the e-vector, but rather by a slight fluctuation in light intensity caused by the polarizer rotation, we projected an unpolarized light stimulus through the depolarizer beneath the rotating polarizer (see Materials and Methods). Under the unpolarized light stimulus, the bees did not show any clear movements coincident with the polarizer rotation (Fig. 4A). Furthermore, no detectable peak at 0.1 Hz was noted in the averaged PS, and none of the five experimental bees demonstrated the highest peak at 0.01 Hz (Fig. 4B). Only one bee showed a small PS peak at 0.01 Hz, which was not significantly different from that under the static e-vector stimulus (*p* = 0.4885, Fisher’s exact test). These results indicate that the abdominal periodic movements were elicited by the rotation of the polarized e-vector orientation.

**Fig. 4.**
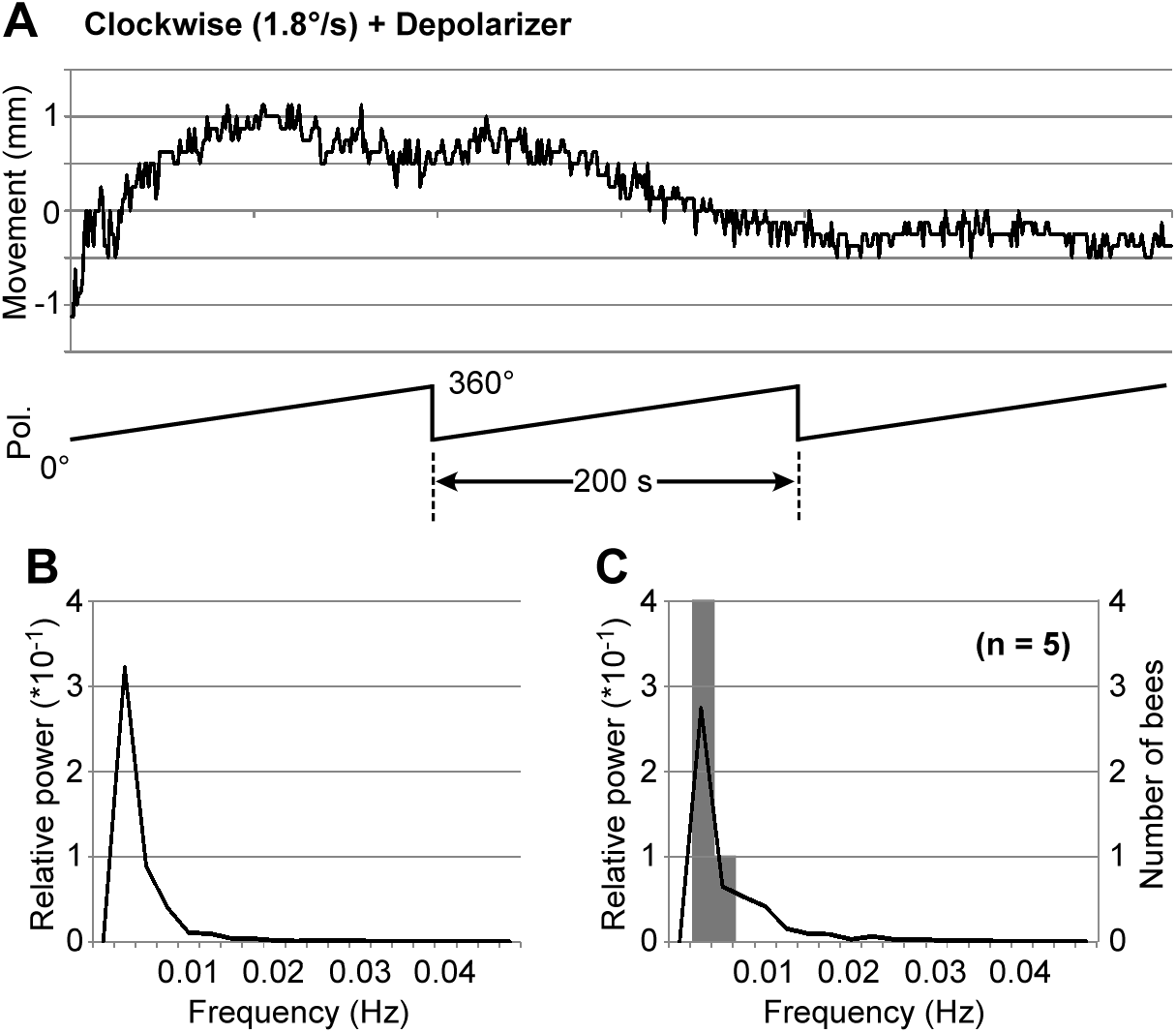
Abdominal movements under the depolarized light stimulus. **A**. An example of the bee’s abdominal trajectory. A UV-transmitted depolarizer was put just below the rotating polarizer (1.8 ° s^-1^). The lower trace (Pol.) indicates the e-vector orientation of the polarizer with respect to the bee’s body axis. **B**. The power spectrum of the abdominal trajectory shown in A. **C**. Averaged power spectrum (black line) and the histogram of the maximum peak in each power spectrum (gray bars) are shown (N = 5).

We also determined the relationship between a tethered bee’s abdominal location and its flying behavior (see Fig. S1). Simultaneous recordings of the abdominal images and the yaw torque of a flying tethered bee showed a strong negative correlation, i.e., the bee’s abdominal tip moved right when the bee turned left and *vice versa*. Therefore, a bee’s periodic abdominal movement under the rotating e-vector stimulus could be considered a kind of polarotaxis to adjust the flying direction to a certain e-vector orientation.

### Polarotaxis under the different speeds of the stimulus

Next, we observed polarotaxis of the tethered bees under CW rotating e-vector stimulus at twice the speed (3.6 ° s^-1^) or 2-times slower the speed (0.9 ° s^-1^) to confirm that the periodicity in the abdominal movement (Fig. 2, 3) was not elicited by internal rhythm but by external polarized light stimuli. Under the faster stimulus, some bees still showed right-and-left abdominal movements synchronized to the stimulus rotation (Fig. 5A). However, in contrast to the 1.8 ° s^-1^ stimulus, the PS of the abdominal trajectory showed only a small peak at a stimulus frequency of 0.2 Hz (Fig. 5B). Moreover, in the averaged PS of all 14 experimental bees, a small, but detectable, peak at 0.02 Hz and the highest peak at 0.0025 Hz were noted (Fig. 5C). The number of bees showing the peak at 0.02 Hz in the PS was significantly different from experiencing the static or 1.8 ° s^-1^ stimulus (7 of 14 bees for 3.6 ° s^-1^ and none of the 21 bees for static and 1.8° s^-1^ stimulus; *p* = 0.0005, Fisher’s exact test), although only one of the 14 experimental bees showed the highest peak at 0.02 Hz (Fig. 5C). These results indicated that the bees exhibited weak polarotactic behavior to the fast rotating e-vector stimulus.

**Fig. 5.**
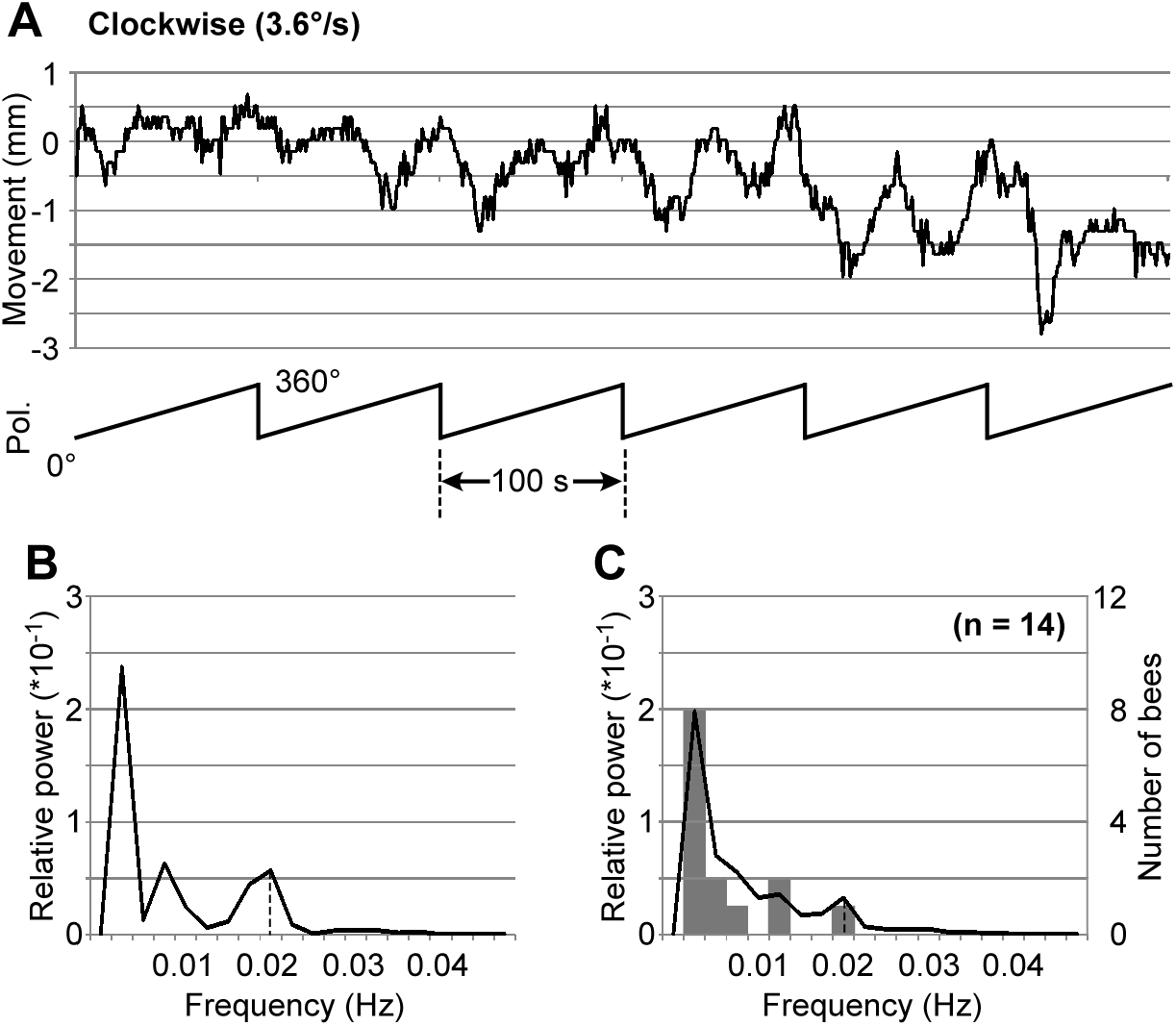
Abdominal movements under the rotating polarized light stimulus (3.6 ° s^-1^). **A**. An example of a bee’s abdominal trajectory. The lower trace (Pol.) indicates the e-vector orientation of the polarizer with respect to the bee’s body axis. **B**. The power spectrum of the abdominal trajectory shown in A. **C**. Averaged power spectrum (black line) and the histogram of the maximum peak in each power spectrum (gray bars) are shown (N = 14). Dashed lines indicate the peaks at the stimulus rotation frequency (0.02 Hz).

Under the slower rotating stimulus, the tethered bees showed clear right-and-left abdominal movements, the PS of which had the highest peak at the stimulus frequency of 0.005 Hz (Fig. 6A, B). Four of the 10 experimental bees exhibited the highest peak at 0.005 Hz in each PS of the abdominal trajectory (Fig. 6C), whereas only one of the 21 bees did so under the 1.8 ° s^-1^ stimulus, which was significantly lower (*p* = 0.0274, Fisher’s exact test). This result indicated that the bees also responded to a slow stimulus. However, we could not detect a 0.005 Hz peak in the averaged PS, although the power at 0.005 Hz was relatively high compared with that under other stimulus conditions (Fig. 6C); this could have occurred because the peak could not be clearly separated from the peak at 0.0025 Hz owing to data interference from unresponsive bees (see Figs 3B, 4C). Probably for similar reasons, the number of bees demonstrated the highest peak at 0.005 Hz was not significantly different from that of the static e-vector stimulus (4 of the 21 bees; *p* = 0.3809, Fisher’s exact test).

**Fig. 6.**
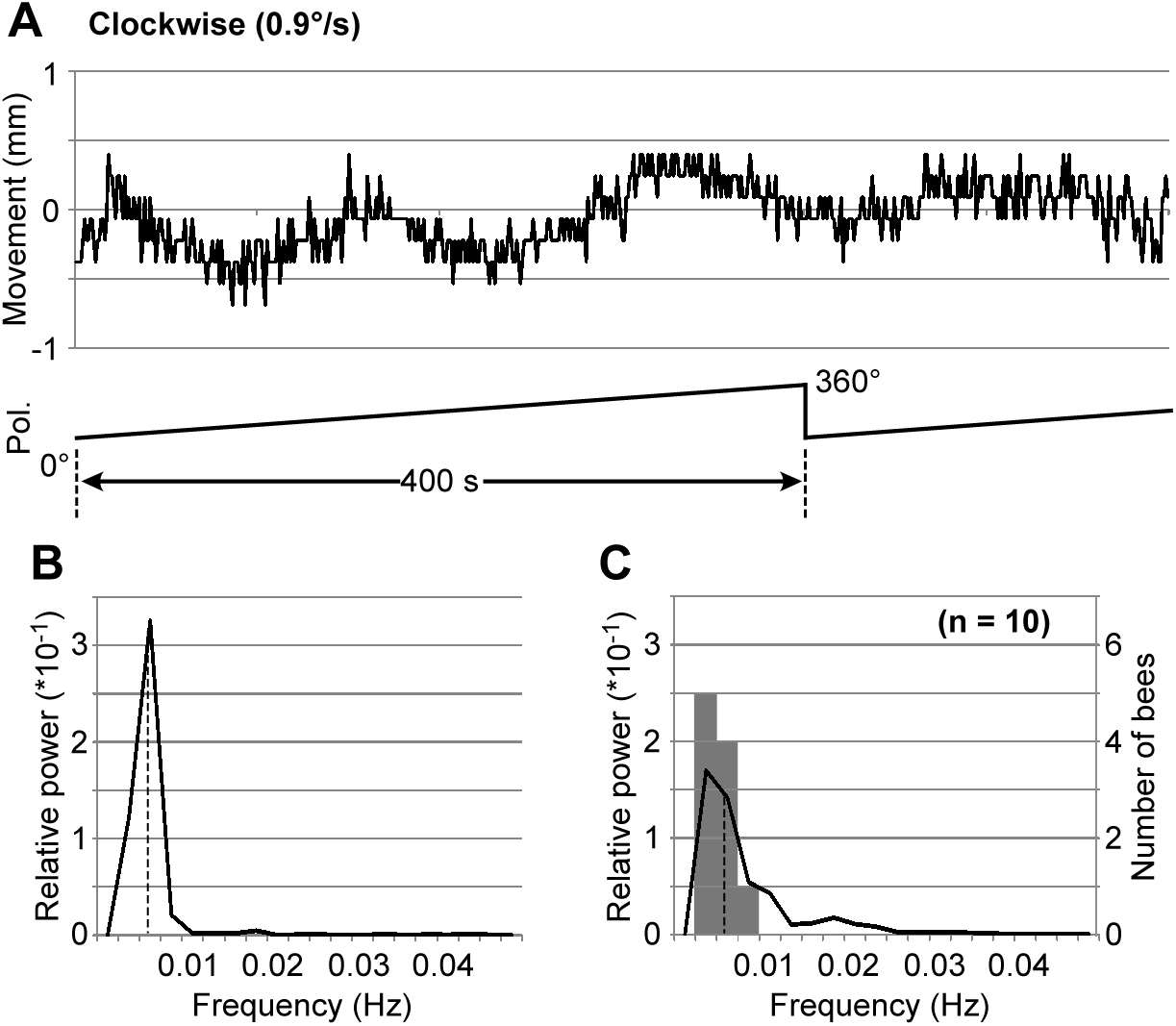
Abdominal movements under the rotating polarized light stimulus (0.9 ° s^-1^). **A**. An example of a bee’s abdominal trajectory. The lower trace (Pol.) indicates the e-vector orientation of the polarizer with respect to the bee’s body axis. **B**. The power spectrum of the abdominal trajectory shown in A. **C**. Averaged power spectrum (black line) and the histogram of the maximum peak in each power spectrum (gray bars) are shown (N = 10). Dashed lines indicate the peaks at the stimulus rotation frequency (0.005 Hz).

### Selective stimulation of eye regions

Polarization vision in insects is known to be mediated by the DRA of the compound eye. To confirm the sensory input area for polarotaxis in the eye, we covered a part of each compound eye and restricted the area receiving light stimulation to the DRA (Fig. 7D, E). The bees whose DRA was covered did not show polarotactic abdominal movement even under the 1.8 ° s^-1^ rotating polarized light stimulus to which intact bees responded (Fig. 7A), and no clear peak was noted at the stimulus frequency of 0.01 Hz in the power spectrum (Fig. 7B). The averaged power spectrum of all eight experimental bees did not exhibit a peak at 0.01 Hz (Fig. 7C), indicating that the bees with covered DRA lost the ability to orient to certain e-vectors. Similar to the response of intact bees to a static stimulus, none of the eight bees displayed the highest peak at 0.01 Hz (Fig. 7C, see also Fig. 3B), and their response was not significantly different (*p* = 1, Fisher’s exact test). Conversely, the number of bees showing the highest peak at 0.01 Hz was also not significantly different than that of the intact bees under the CW stimulus (see Figs 3A, 7C; *p* = 0.1421, Fisher’s exact test), probably owing to the small number of experimental bees used.

**Fig. 7.**
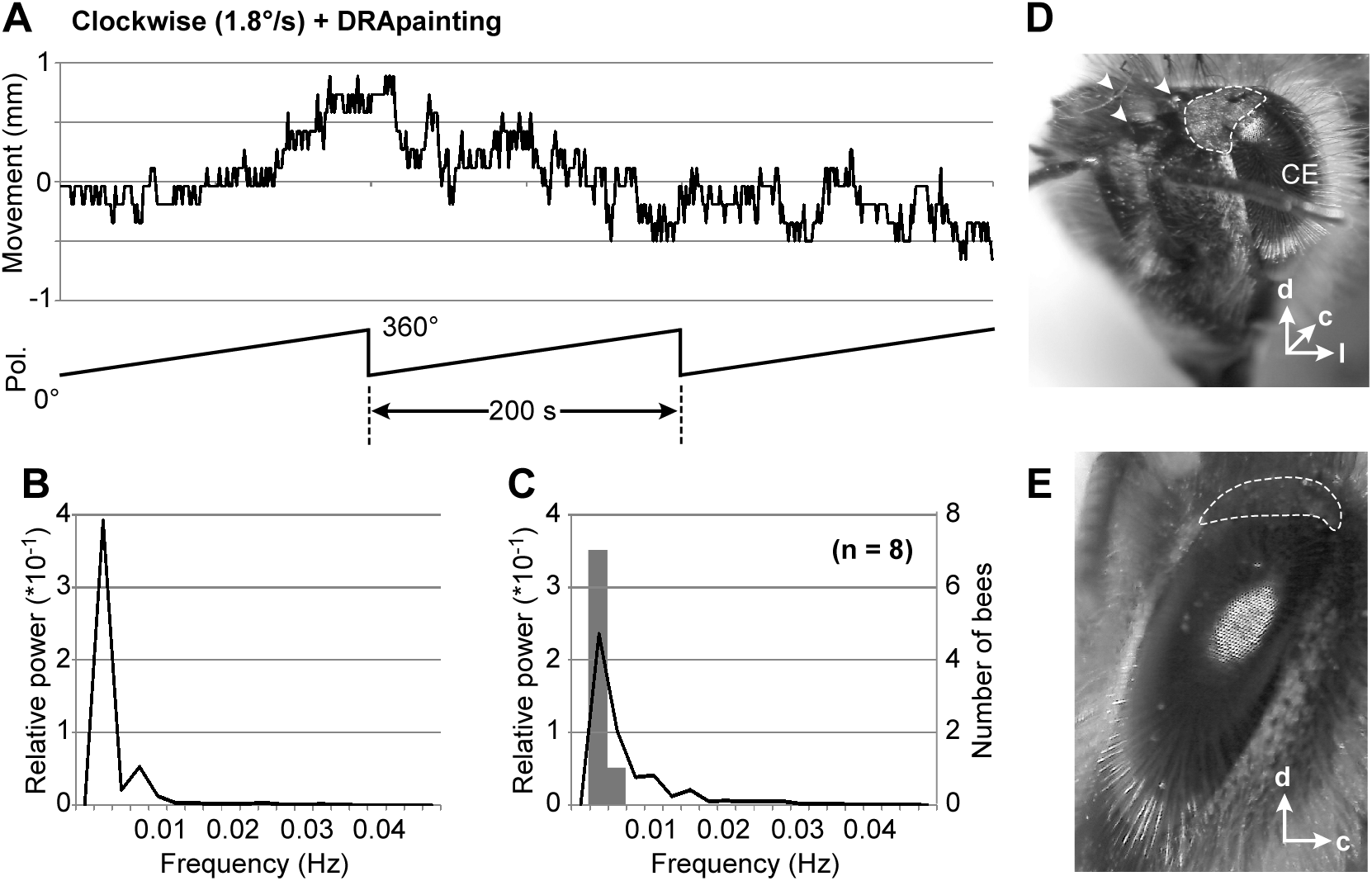
Abdominal movements of the DRA-covered bees under the rotating polarized light stimulus (1.8 ° s^-1^). **A**. An example of a bee’s abdominal trajectory. The lower trace (Pol.) indicates the e-vector orientation of the polarizer with respect to the bee’s body axis. **B**. The power spectrum of the abdominal trajectory shown in **A**. **C**. Averaged power spectrum (black line) and the histogram of the maximum peak in each power spectrum (gray bars) are shown (N = 8). **D**. Head of the bee after its DRAs were painted. The area surrounded by the dashed line was painted. Arrow heads indicate the ocelli. CE: compound eye. **E**. Lateral view of the compound eye of the bee shown in **D**.

### Preferred e-vector orientation

We assessed the PEOs of the 21 bees that showed polarotaxis under the 1.8 ° s^-1^ CW stimulus. The PEO of each bee varied from 0 to 180 ° (Fig. 8A) However, more than half of the bees (14 of 21) showed PEOs between 120 to 180 ° and the distribution was not significantly random (*p* = 0.009, Rayleigh test). To confirm whether each bee had a specific PEO, we compared the PEO to the CW and CCW stimuli in the same bee. The angular differences in PEOs between CW and CCW stimuli of 7 bees, which showed clear polarotaxis under both conditions, are shown in Fig. 8B. The distribution of the angular differences was not significantly random (*p* = 0.01, Rayleigh test), but was concentrated around 0 ° (*p* = 0.026, *V*-test), suggesting that each bee had a certain PEO and adjusted the flight direction by aligning to a particular e-vector angle.

**Fig. 8.**
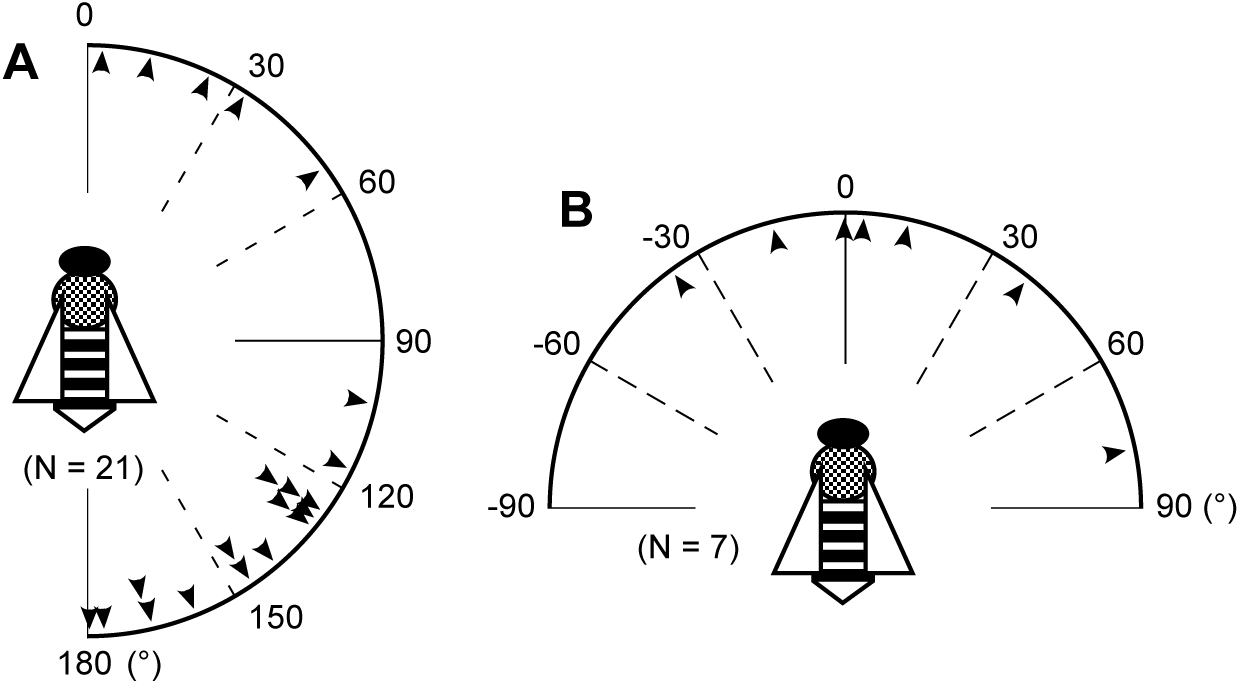
Preferred e-vector orientations (PEOs) of the bees caught at the hive entrance. **A**. PEOs of the bees that showed polarotaxis under clockwise rotating stimulus (1.8 ° s^-1^) with respect to the bee’s body axis. The distribution was significantly random (*p* > 0.1, Rao’s spacing test). **B**. Angular differences in the PEOs of each bee between clockwise and counterclockwise stimulus. The distribution was not significantly random (*p* < 0.05, Rao’s spacing test), but concentrated around 0 ° (*p* = 0.026, *V*-test).

## DISCUSSION

### Behavioral response to the polarized light stimulus in the honeybee

It is well known that honeybees use skylight polarization to detect their intended travel direction. It was first described by von Frisch (1967) through a series of sophisticated behavioral studies on the waggle dance. Thereafter, the waggle dance orientations of the nest-returning bees from a certain feeder have been intensively studied. These studies were conducted under a patch of polarized light stimulus or part of the sky and an internal representation of the celestial e-vector map has been proposed (Rossel and Wehner, 1982, 1986, 1987; Wehner, 1997). These systematic studies have focused on modification of the waggle dance orientation and not on how the bees perceive polarized light from the sky en route to/from the nest. More recently, direct observation of polarized light detection in flying bees has been performed using a four-armed tunnel maze with a polarizer on top (Kraft et al., 2011). In this experiment, it was revealed that bees choose their flying direction based on zenithal polarized light information. Moreover, it has also been demonstrated that bees memorize the e-vector orientations experienced during their foraging flight and use that memory for the subsequent waggle dances (Evangelista et al., 2014). In the present study, we directly showed that bees tended to orient to the certain e-vector angles during their flight under the tethered condition, i.e., they referred polarized light information to control their flight direction. The fact that fewer bees responded to the fast stimulus (3.6 ° s^-1^, Fig. 5) than the slow stimuli (0.9 ° s^-1^ and 1.8 ° s^-1^, Fig. 2, 3, 6) is also indicative of the use of e-vector orientation as a global cue for orientation. Probably, they did not refer to the e-vector when it quickly changed because they did not expect such a situation, except when they quickly changed their flight direction.

The polarotactic behaviors were not observed when the bee’s DRA was blinded (Fig. 7). It is well known that detection of skylight polarization in insects is mediated by ommatidia in the DRA (for review: Labhart and Meyer, 1999; Wehner and Labhart, 2006). In honeybees, UV-sensitive photoreceptors of the ommatidia in DRA are highly polarization-sensitive, and their receptive field covers a large part of the celestial hemisphere, which is suitable for observing the sky (Labhart, 1980, Wehner and Strasser, 1985). Behaviorally, it has also been demonstrated that covering the DRA impaired correct coding of food orientation by the waggle dance orientation (Wehner and Strasser, 1985). These results clearly show that bees utilize polarized light detected by the ommatidia in the DRA for orientation.

### Polarotaxis in insects

Polarotactic behavior in insects has been demonstrated in several species. Obviously, orientation to a certain e-vector direction is a common occurrence among insect species that utilize skylight polarization for navigation. Classically, it has been tested using a treadmill device in the cricket *Gryllus campestris* (Brunner and Labhart, 1987) and the fly *Musca domestica* (von Philipsborn and Labhart, 1990). Using such a device, the insect was tethered on an air-suspended ball and its walking trajectory could be monitored through the rotation of the ball. In these species, the insect on the ball showed clear polarotactic right-and-left turns when the e-vector of the zenithal polarized light stimulus was slowly rotated, as we showed in this study in flying honeybees. This kind of behavior does not merely demonstrate they have polarization vision but also allowed us to clarify fundamental properties of insect polarization vision, e.g., perception though the DRA in the compound eye (Brunner and Labhart, 1987), monochromatic spectral sensitivity (Herzmann and Labahrt, 1989; von Philipsborn and Labhart, 1990), and sensitivity to the degree of polarization (Henze and Labhart, 2007).

Orientation to the polarized light has been investigated in tethered flying insects as well. In the locust *Schistocerca gregaria*, direct monitoring of yaw-torque responses showed clear polarotactic right-and-left turns to rotating polarized light (Mappes and Homberg, 2004). In tethered monarch butterflies *Danaus plexippus*, measuring flight orientations using an optical encoder revealed that their flight orientation under natural skylight was clearly affected by a dorsally presented polarization filter (Reppert et al., 2004; but see also Stalleicken et al., 2005). Similar orientation to polarized skylight has also been demonstrated in *Drosophila* (Weir and Dickinson, 2012). In this study, a fly was magnetically tethered in the arena and its flight heading was recorded from above by an infrared camera. A potential problem in investigating polarization vision in tethered flying insects is that sometimes the tethering apparatus, including the torque meter or other recording devices, interrupt a part of the visual field of the tested animal. In the present experiments, we succeeded in evaluating the bee’s polarotactic flight steering by simply monitoring the horizontal position of the abdominal tip that was strongly anti-correlated with the torque generated by the bee (see Fig. S1). Using these methods, the entire visual field of the animal remained open; therefore, it had an advantage for investigating the animal’s responses under various stimulus conditions.

### Preferred e-vector orientation

The PEO distribution has been reported in several species. In crickets and flies, a weak preference to an e-vector orientation perpendicular to their body axis was demonstrated, although the reason of this behavior was not clear (Brunner and Labhart, 1987; von Philipsborn and Labhart, 1990). On the other hand, in laboratory-reared locusts, the PEOs were randomly distributed and they did not show any directional preferences (Mappes and Homberg, 2004). In the present study, the bees showed a significantly higher preference between 120 and 180 ° (Fig. 8A). In each bee, the PEOs under CW and CCW stimulus were quite similar (Fig. 8B), and this suggested that each bee has its own PEO and used it not only as a reference for maintaining straight flight but also to deduce its heading orientation.

Considering that central place foragers, such as honeybees, have to change their navigational directions depending on the currently available food locations, their PEOs would reflect their previous foraging experiences. In the present study, we collected the bees with a pollen load at the hive entrance; therefore, all experimental forager bees were returners. Consequently, we could no longer assess their feeding locations when we measured their flight responses in the laboratory. Moreover, their path-integration vector should be reset to a zero-state in such a situation (Sommar et al., 2008), and they might not have had a strong motivation to use polarized light cues for navigation. To further clarify the role of polarization vision in flying foragers, testing the PEOs in the bees in different navigational states will be crucial.

## Abbreviations

CW: clockwise
CCW: counterclockwise
DRA: dorsal rim area
PEO: preferred e-vector orientation

## Acknowledgements

The authors are deeply grateful to Dr. M. V. Srinivasan and Dr. T. Luu for helpful advice regarding the construction of the flight simulator. We also thank Dr. Y. Hasegawa, Dr. H. Ikeno, and Ms. H. Onishi for their help measuring the torque of flying bees.

## Competing interests

No competing interests declared.

## Funding

This work was supported by KAKENHI from the Japan Society for Promotion of Science (15KT0106, 16K07439, and 17H05975 to MS; 16K07442 to RO) and Bilateral Joint Research Project with Australian Research Council from Japan Society for Promotion of Science to MS.

